# Small molecules enhance the potency of natural antimicrobial peptides

**DOI:** 10.1101/2021.07.06.451332

**Authors:** Valeria Losasso, Khushbu Agarwal, Morris Waskar, Amitabha Majumdar, Jason Crain, Martyn Winn, Michael Hoptroff

**Affiliations:** Science and Technology Facilities Council, Daresbury Laboratory, Sci-Tech Daresbury, Daresbury, WA4 4FS; Unilever Research and Development, Bangalore 560066, India; IBM Research Europe, Hartree Centre, Daresbury WA4 4AD, U.K; Unilever Research and Development, Port Sunlight, UK, CH63 3JW; Department of Biochemistry, University of Oxford, Oxford OX1 3QU, U.K

**Author notes:** For correspondence: Michael Hoptroff.

**Keywords:** antimicrobial peptide (AMP), computer modelling, membrane bilayer, *Staphylococcus aureus* (*S. aureus*), skin

## Abstract

The skin-associated microbiome plays an important role in general well-being and in a variety of treatable conditions. In this regard, endogenous antimicrobial peptides have a role in controlling the microbial population. We demonstrate here that certain small molecular species can amplify the potency of naturally-occurring antimicrobial peptides. For example, Niacinamide is a vitamin B3 analogue naturally found in foods and widely used in topical skin care products, and here we have investigated its cooperativity with the human antimicrobial peptide LL37 on the bacterium Staphylococcus aureus. We have also studied two other structurally related B3 analogs. We observed a clear synergistic effect of niacinamide and, to some extent, methyl niacinamide, whereas isonicotinamide showed no significant cooperativity with LL37. Adaptively-biased molecular dynamics simulations revealed that the analogs partition into the head group region of an anionic bilayer used to mimic the bacterial membrane. The observed effects on the physical properties of the membrane are well correlated with experimental activity. In contrast, the analogs have little effect on zwitterionic bilayers which mimic a mammalian membrane. We conclude that these vitamin B3 analogues can potentiate the activity of host peptides by modulating the physical properties of the bacterial membrane, and to a lesser extent through direct interactions with the peptide. The level of cooperativity is strongly dependent on the detailed chemistry of the additive, suggesting an opportunity to fine-tune the behaviour of host peptides.

## Introduction

It is increasingly recognised that consumer wellbeing issues such as body malodour [1] and dandruff [2] and conditions such as atopic dermatitis [3] are commonly associated with imbalances in the skin microbiome. There is an emerging consensus that the Staphylococcal population of the skin microbiome, reflected in the ratios of *S.hominis* to *S.epidermidis* and *S.capitis* to *S.epidermidis* are implicated in axillary malodour and in scalp health, respectively. There is also increasing recognition of the importance of *S.aureus* colonisation of human skin in atopic dermatitis [3, 4].

*Staphylococcus aureus* colonisation of the human population is widespread with approximately 30% of all humans carrying the organisms as a benign microbial inhabitant of the internal nasal cavity [5, 6]. However, although a benign inhabitant of the nose, colonisation by *S. aureus* of the exposed skin of the face or body is commonly associated with negative pathologies including atopic dermatitis where the mean relative abundance of *S. aureus* has been reported as increasing to 65% during an atopic flare event compared to 1.1% in healthy controls [3].

The human body has evolved ways to modulate the human-associated microbiome through mechanisms including the production of antimicrobial peptides (AMPs), a family of small endogenously produced compounds released through sweat and sebaceous secretions [7]. They are a primitive form of defense mechanism and part of the innate immune system. AMPs adsorb to bacterial cell membranes by electrostatic attraction due to the presence of acidic phospholipids, which confer a net negative charge to the membrane, leading to AMP aggregation and integration, and local membrane thinning [8]. AMPs can target other structures or processes of microbes, such as internal organelles or they may inhibit enzyme activity or macromolecule synthesis [9]. Keratinocytes – forming a large proportion of normal, healthy skin cells, produce antimicrobial peptides, such as the human cathelicidin LL-37, beta-defensins 2 and 3 and dermcidin, which contribute to the skin’s ability to deter the overgrowth of undesirable micro-organisms [10]. AMPs are expressed by keratinocytes either constitutively or are upregulated in response to microbial stimuli. Commensal and pathogenic staphylococci have been shown to activate different pathways in human keratinocytes, and commensals are able to amplify the innate immune response of keratinocytes to pathogens [11].

In the skin, the two best characterised families of AMPs are the defensins and cathelicidins. Defensins are packed in lamellar bodies within keratinocytes and release to the cell surface [12]. Human beta defensin-2 (HBD2) is active against gram negative bacteria, while Human beta defensin-3 (HBD3) kills both gram-negative and gram-positive bacteria such as *S. aureus* [13]. Within the cathelicidin group, LL37 has a broad activity spectrum and is reported to be effective against gram-positive and gram-negative bacteria, and viruses [12]. Other AMPs that also play a role in skin defence against pathogens include Psoriasin, which shows activity against *E. coli,* and RNase7, which has activity against *S. aureus* as well as gram negative bacteria [12]. In addition to their antimicrobial role, AMPs also function as immune-modulators that could help supplement skins defences.

Discovering how AMPs exert their antimicrobial effect and translating this insight into consumer products which work in partnership with natural defence peptides is important when identifying innovative sustainable technologies for consumers. The observed target specificity suggests that AMPs are sensitive to the composition of the lipid membranes. This raises the possibility that AMP activity may be modulated by influencing lipid composition of the microbial target, rather than the more usual route of re-designing the AMP.

Several mechanisms may exist by which small molecules may potentiate the activity of endogenous antimicrobial peptides such as LL37. Such potentiators could display a biological mechanism, for example by increasing the biological expression of functional AMPs or by increasing their conversion from less antimicrobial pro-forms. Alternatively, they may participate directly in the antimicrobial activity, for example by interacting with the AMP at a structural level or by an indirect physical mechanism such as destabilisation of the bacterial membrane and enhancing the action of the AMP. Although it has been reported previously that small molecules, like niacinamide that are not well-known as direct antimicrobials, nonetheless give hygiene benefits by enhancing the expression level of AMPs in human tissue [14]. However, no work to our knowledge has investigated the potentiation of antimicrobial activity of AMPs through such small molecules like niacinamide.

This paper therefore seeks to explore the hypothesis that small molecules may amplify the potency of naturally occurring AMPs through mechanisms of physical interaction, in addition to the biological mechanisms previously reported [14] In order to properly and precisely investigate such physical mechanisms, this investigation is focussed on in-vitro models of AMP and potentiator interaction with bacterial targets and has deliberately excluded the investigation through the use of viable human cell models.

Niacinamide is naturally found in foods, and is widely used as a cosmetic skin care ingredient which has been also shown to mitigate against infection in mice [15] and to enhance AMPs in gut epithelial cells, in neutrophils [16], and in skin cells [14].

Here, we use computer simulations to investigate how AMPs and candidate potentiator molecules interact with bacterial membranes at the molecular scale and explore correlations with microbial growth in a well-controlled single species in-vitro model of microbial growth inhibition by LL37. To demonstrate the specificity of the interaction to bacterial membranes, similar work was completed on model non-bacterial membranes.

Mammalian cytoplasmic membranes contain mainly phospatidylcholine, phosphatidyletanolamine (PE) and cholesterol. The main components of bacterial membranes are, on the other hand, PE, phosphatidylglycerol and cardiolipin (CL) [17]. Notably, all bacterial membranes have anionic lipids in their composition which can be PG and/or CL. The presence of these anionic lipids explains the selective toxicity of AMPs against bacteria but not against mammalian cells [18],

Simple phosphatidylcholine (POPC) and phosphatidylglycerol (POPG) models representing neutral mammalian membranes and negatively charged bacterial membranes are routinely used in studies to easily represent differences between the two systems (see for example [19–21] and our previous work on AMPs [22]. Therefore, we chose to use POPC and POPG, consistent with a recent simulation study on LL-37 [23].

## Results

### Activity against S. aureus

The synergy between LL37 and each of the additives against a cell culture of *S. aureus* was evaluated using modified microdilution assays, as described in the Experimental Procedures. The reactions were performed in a low salt buffer as high concentrations of salts are known to be inhibitory to AMP activity. Figure 2 shows the bacterial recovery rate averaged over four replicates taken from two independent experiments. While niacinamide has no effect on the recovery rate by itself, it increases the measured activity of LL37 compared to LL37 alone. A similar but smaller effect is seen for methyl nicotinamide. In the case of isonicotinamide, the effect is not significant at a 5% confidence threshold. Given the chemical similarity of the three additives, these differences are striking but reproducible (see TableS1 for individual results).

### Interaction of additive molecules with model membranes

In order to explore the assay results at the molecular scale, we performed computer simulations of the relevant molecules in model membranes. We began by considering the interaction of niacinamide and its analogues on their own with a lipid bilayer, in order to provide a reference point for later simulations with AMP molecules included. We chose POPC and POPG as simple model membranes for human and bacterial cells, respectively, a model consistent with a recent simulation study on LL37 [23].

The association of molecules with lipid bilayers and subsequent insertion kinetics may encounter significant energy barriers which present challenges for conventional molecular dynamics methods to sample correctly. Here we use adaptive-biasing-force simulations to overcome barriers in the free energy landscape. Integration of the average force along a chosen reaction coordinate gives a measure of the free energy of insertion for these molecules through the potential of mean force (PMF), see Figure 3. Insertion is generally unfavourable for niacinamide and its derivatives, except around the headgroup region, which is consistent with negative logP values determined experimentally (e.g. as obtained from PubChem entries). Notably, the niacinamide preferential localization is in contrast with the behaviour reported in PMF studies for a similar molecule, thymol, which shows instead a clear propensity to insert into the hydrophobic interior of a DPPC model membrane [24]. This can be explained by the higher polarity of niacinamide due to the presence of both the pyridine nitrogen and the amide group as substituents.

All molecules show a local minimum in the free energy between 2 and 5 Å below the membrane surface. For each molecule the process of reaching the surface is more favourable for negatively charged POPG than for POPC, as a result of the polarity of the substituent groups. Thus, the free energy surface for niacinamide and n-methylnicotinamide is relatively flat from 3Å on either side of the POPG membrane surface.

Unbiased simulations were used to confirm the PMF results and provide more detail on molecular interactions. Additive molecules were initialised at a depth of 5 Å in the membrane, in the region of the free energy minima shown by the PMF curves. We examined all three additives, and give example results for the case of niacinamide.

An analysis of the membrane thickness and the area per lipid across the simulated patch shows that niacinamide induces thinning and stretching of the POPG membrane (Fig. S1a-b), however has little effect on POPC membranes. The deuterium order parameter (S_CD_) order parameter shows that it induces disorder along the lipid tails for a POPG membrane, while the effect is not observed for POPC (Fig. S1c). The lipid tail disorder is also shown by an increase in lipid tilt angle with respect to the membrane normal when niacinamide is simulated in combination with POPG (Fig. S1d). Finally, niacinamide increases POPG membrane fluidity, as indicated by the ensemble average of time-averaged MSDs (TAMSD) as a function of the measurement time (Fig. S1e).

Focussing on the POPG membrane, a collective analysis of niacinamide together with its derivatives shows that they affect membrane properties in the order niacinamide > n-methylnicotinamide > isonicotinamide (Figure 4). Following this order, all three additives decrease POPG thickness (Fig. 4a) and increase area per lipid (Fig. 4b), although niacinamide displays some variability between the two simulation replicas resulting in a bimodal distribution. The three molecules all decrease the SCD order parameter, with small differences among them (Fig. 4c). However, the lipid tilt angle shows a significant increase in tail disorder in the presence of niacinamide compared to its derivatives and to pure POPG (Fig 4d).

Fig. 4E shows the ensemble average (over all lipid molecules) of the time averaged MSDs for POPG as a function of the measurement time. These values suggest that the lipid mobility is also higher in the presence of niacinamide. This quantity is indeed linked to lateral diffusion coefficients for lipids [25] and it shows the effect of the presence of the additives in the first few hundreds of nanoseconds of simulation; the long time behaviour, with all systems close to a value of ~0.4 Å^2^/ns although not fully converged, is in agreement with experimental values reported for POPG lateral diffusion coefficients [26]. These results are consistent with a mechanism where additives perturb the membrane on a short timescale (< 1 us), which is however sufficient for AMPs to exert their action as demonstrated in Fig. 5a.

In summary, niacinamide and its analogs partition into the head group region of POPC or POPG bilayers. All three additives have a clear effect on the physical properties of the POPG membrane, although the extent varies, while the effect on POPC membranes is less clear. These results suggest that a suitable concentration of additives could alter how anionic membranes interact with AMPs.

### Stability and orientation of LL37 in model membranes

We next considered the interaction of a single AMP, as exemplified by LL-37, with the POPC and POPG model membranes in the absence of additive molecules. We analysed the ability of the peptide to enter and deform the lipid bilayers, as well as the conformational changes of the peptide itself.

Starting from an initial position above the membrane surface, LL-37 locates in the headgroup region of POPG membranes, whilst it does not bind POPC membranes and remains in aqueous solution, as shown by the distribution of distances to the membrane surface (Figure 5a). Consistently with all helical AMPs [27], it preserves most of its secondary structure when bound to POPG (Fig. 5b), whilst it shows partial disruption of its helical configuration in the presence of POPC (Fig. 5b) where the water environment contributes to the unfolding of the region between residues 8-15 (Fig. 5c).

In binding POPG, LL-37 also affects the physical properties of the membrane (Fig. S2). In particular, it reduces its average thickness (Fig. S2-a) and increases its disorder and fluidity (Fig. S2 c-e), although does not affect significantly its area per lipid (Fig. S2-b). In contrast, there is no significant effect on POPC membranes.

### Interaction of additive molecules with LL37

We next considered direct interactions of additive molecules with LL-37. Given the failure of LL-37 to insert into the POPC membrane, we focus here on the observations for the POPG membrane system. Simulation of model POPG membranes with both additives and LL-37 included revealed transient contacts (Table S1, Figure 6a). However, we did not observe any stable complexes between the additive molecules and LL-37 in membrane.

The lack of contacts may be due to poor sampling in the MD simulation, and so we also considered additives and LL-37 in solution. To mimic different environments and their effect on electrostatic interactions, we considered three solvents: water, methanol and octanol.

In water, all niacinamide derivatives show a few transient contacts with LL-37 (Figure 6b). In contrast, in hydrophobic environments, we observe a differentiated effect: while in the membrane the largest number of contacts were observed for n-methylnicotinamide (Figure 6a), niacinamide shows more contacts in hydrophobic solvents (methanol and octanol, Figure 6c-d). Interestingly, isonicotinamide showed few contacts in all solvents, despite being an isomer of niacinamide. Isomerisation leads to a charge redistribution in the conjugated system (Fig. S3).

We also analysed the formation of hydrogen bonds between additives and LL-37 in the four different contexts, to investigate the chemical features underlying the differences between niacinamide and its analogues, in particular its isomer isonicotinamide. We did not observe stable hydrogen bonds in the membrane systems or in water (Table S2), therefore we used hydrophobic solvents to mimic the membrane environment while allowing for a faster diffusion.

In methanol, we start observing more significant hydrogen bonds between the amide oxygen of niacinamide and positively charged residues of the central region of LL-37 (Lys12, Lys15, Lys18, Arg19, Arg23), which become less frequent in the case of n-methylnicotinamide and isonicotinamide. However, clearer differences among the three analogs are observed in octanol, where niacinamide shows more overall contacts with respect to its derivatives, especially isonicotinamide. Niacinamide forms stable and sustained contacts between its polar atoms and a small group of residues (Lys12, Lys15, Lys18, Arg19, Gln22, Arg23), which are also found at a lower extent in n- methylnicotinamide but at a much lower extent in isonicotinamide (Table S2). The simultaneous formation of two hydrogen bonds between the same niacinamide molecule and pairs of these residues, enabled by the specific configuration of the amide substituent compared to isonicotinamide, might explain the higher frequency of these contacts in the presence of the first ligand. In particular, niacinamide amide oxygen and ring nitrogen can form simultaneous interactions with Lys12 and Lys15 side chains as well as with Gln22 and Lys18 side chains, respectively. (Figure S4).

### Combined effect

Finally, we analysed the effect on POPG membrane properties of the combined presence of additives and LL37 compared to a control membrane in complex with LL37 only. The results can also be compared with the membrane properties in the presence of the additives only, shown in Figure 4a-d and replotted in Figure 7.

In general, for niacinamide + LL-37 we see that thickness decreases, area per lipid increases, and lipid order (measured by S_CD_ parameter and lipid tilt angle) decreases, compared with the pure membrane or either component individually. These effects can also be seen in the other additives to a lesser extent, and we can again place the analogues in the order niacinamide > n-methylnicotinamide > isonicotinamide (Figure 7). In particular, only niacinamide shows a strongly cooperative effect with LL-37 wherein the small molecule and AMP in combination impact membrane properties to a greater extent than either one individually (Table S2).

## Discussion

Our experimental assays have revealed that some small molecule additives can potentiate the activity of the naturally occurring endogenous skin AMP LL-37 against *Staphylococcus aureus*, an organism closely associated with atopic dermatitis [3–4, 28–29]. Common cosmetic ingredients like niacinamide could potentially work with the innate defenses of the skin and provide enhanced protection against pathogens such as *S. aureus*. This understanding may be expanded to other harmful strains such as MRSA that cause severe disease and need to be controlled. There are other benefits from niacinamide reported in literature, including boosting of antimicrobial peptides [14], and the synergy mechanisms reported here could work along with them. Taken together our data demonstrates how a molecule such as niacinamide, which is not inherently antimicrobial, may enhance hygiene benefits by potentiating the body’s natural defenses by a two-pronged mechanism which includes increasing the number of AMPs [14] and, as reported here, by potentiating their activity against pathogens. Such multifunctional technologies could provide the necessary benefits not just to the host, but also keep undesirable microbes in check.

Using molecular simulations of these molecules and a model membrane, we have discovered two distinct mechanisms by which this synergy can occur. Firstly, the additives alter lipid bilayer properties such as thickness, area per lipid and acyl tail disorder, and may make the membrane more susceptible to AMPs. Secondly, the additives interact directly with the AMP, especially in more hydrophobic environments.

The strength of this effect depends significantly on the lipid type and on the chemistry of the additive. The results are much clearer for the anionic POPG bilayer, which provides a simple model of a negatively charged bacterial membrane. There is also large variation between the three additives studied, even though they are all vitamin B3 analogs.

Niacinamide partitions into the head group region of the bilayer. Although it does not fully permeate, it is still observed to have an effect on membrane physical properties. This is an example where traditional metrics, such as logP, can be misleading. It is also observed to make transient contacts with LL-37 both in solution and in the membrane, suggesting a direct synergistic effect. More stable interactions, including specific hydrogen bonding, are observed in hydrophobic solvents which are often used to represent the membrane interior. Although we do not observe niacinamide partitioning into the interior, this may be relevant in more complex membranes or upon AMP-led perturbation.

Isonicotinamide, although an isomer of niacinamide, is observed experimentally not to have a significant synergy with LL-37. Simulations support this result, in terms of its effect on the physical properties of model membranes and the lack of interaction with LL-37. Although some effects are seen in simulation, these are less clear-cut than for niacinamide and may not be significant physiologically. N-methylnicotinamide presents an interesting case. While most metrics show an intermediate effect, it exhibits a higher number of contacts with LL-37 in the membrane (Fig. 6a). This could imply a slightly different balance between the two proposed modes of action, i.e. destabilisation of the membrane versus direct interaction with LL-37.

Our study has elucidated the mechanisms by which subtle changes in structure influence observed cooperativity between certain small molecules and natural peptides. Further research would be valuable to determine how these insights apply to other small molecules, and to optimise the potentiation of AMPs.

Ultimately, this work enhances our understanding of how the activity of naturally occurring skin AMP’s may be impacted by small molecules and as such opens opportunities for greater understanding of both their efficacy in-situ and modulation by topical treatments..

## Experimental Procedures

We assessed the following series of additives for their ability to act as potentiators of AMPs: niacinamide, isonicotinamide and N-methylnicotinamide (Figure 1). These are all naturally occurring analogs of vitamin B3. As an exemplar AMP, we chose the human peptide LL-37, with sequence LLGDFFRKSK–EKIGKEFKRI–VQRIKDFLRN–LVPRTES.

**Figure 1:**
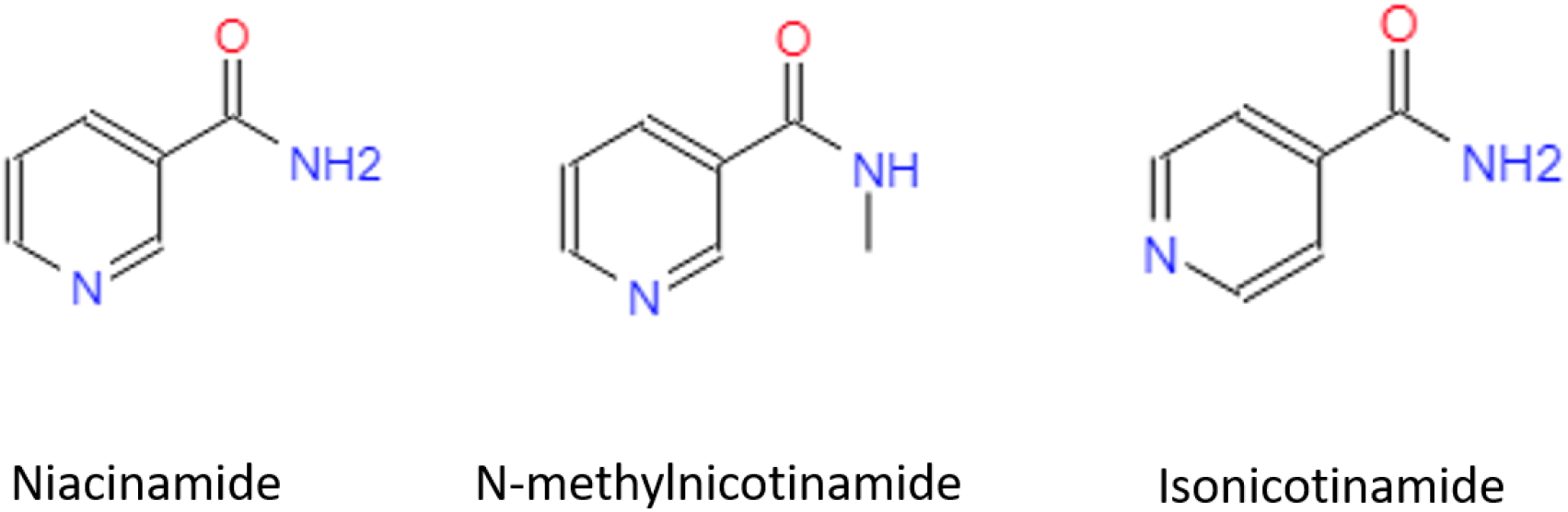
Additive molecules considered.

**Figure 2:**
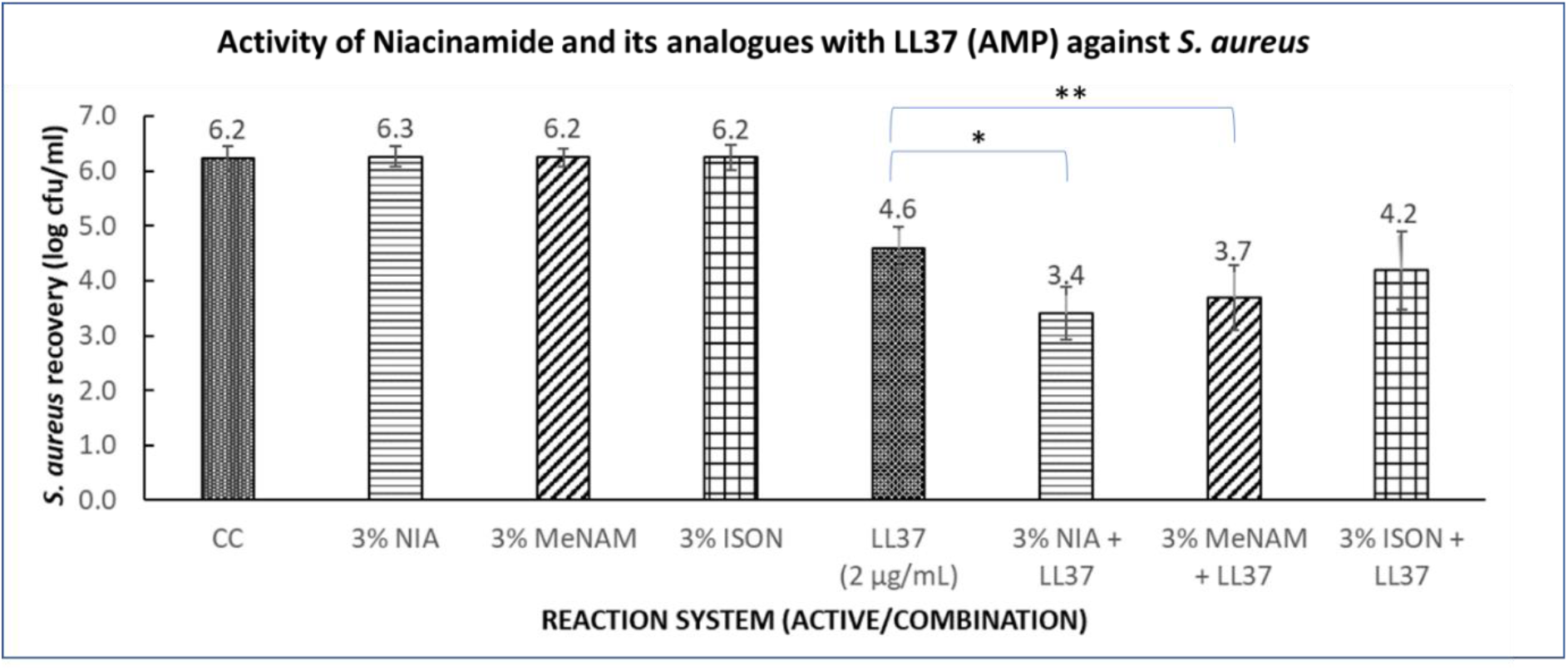
Microdilution Assay of niacinamide and its analogues against *S. aureus*. Niacinamide shows significant amplification of LL37 potency, this synergy is less with methylnicotinamide, while no significant synergy was observed with isonicotinamide. KEY: CC= Culture control (no treatment), NIA= niacinamide, MeNAM=N-methylnicotinamide, ISON= isonicotinamide, LL37= Cathelicidin antimicrobial peptide (AMP), used at 2 microgram per ml concentration. Data is from two independent repeat experiments, each comprising two replicates. Error bars are SE of mean. * and ** are p<0.05 (one tailed t-test)

**Figure 3.**
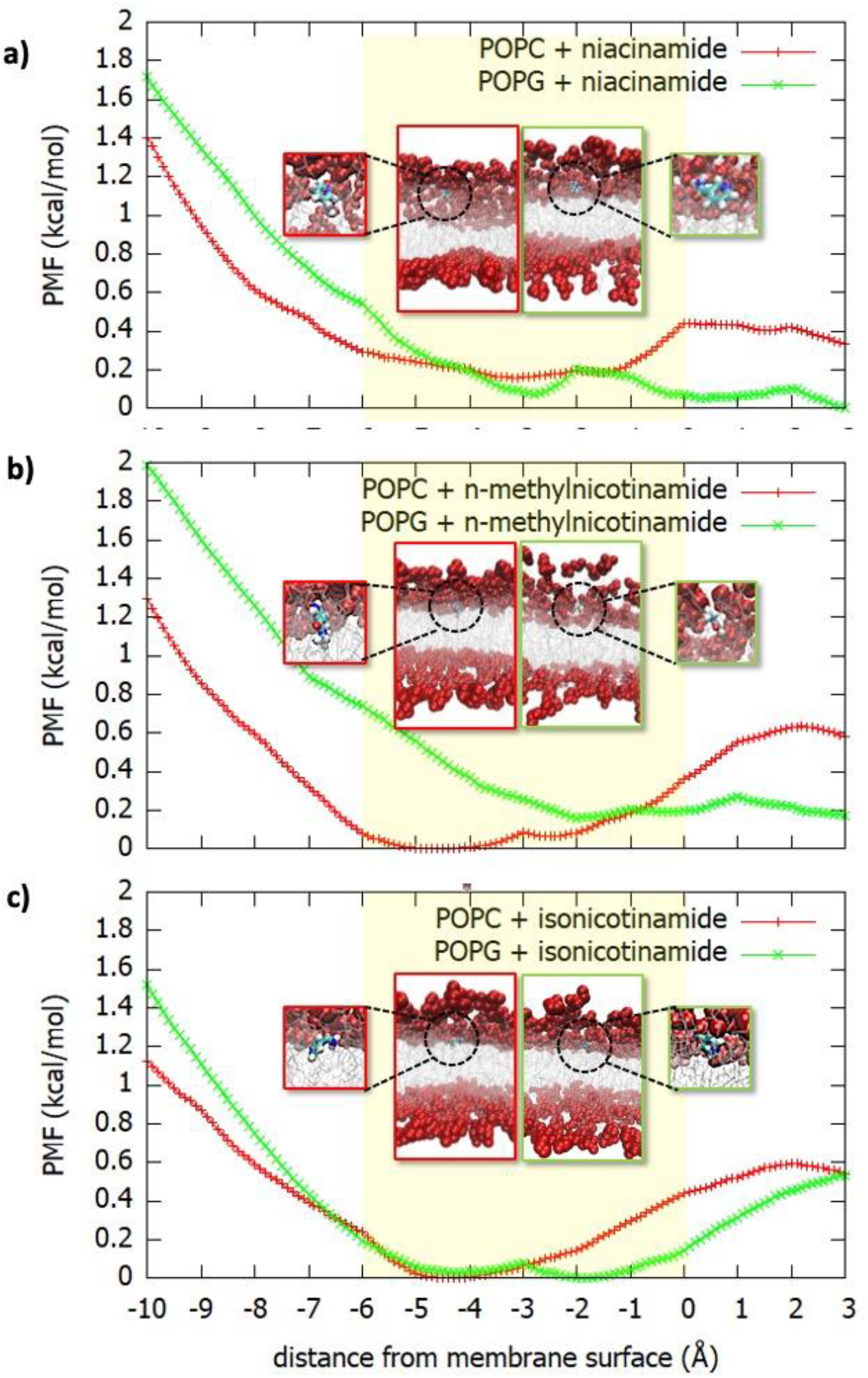
PMF showing the free energy for insertion of a) niacinamide, b) n-methylnicotinamide and c) isonicotinamide in POPC and POPG model membranes. The PMF is given as a function of the distance from the membrane surface, defined as the average z coordinate of upper leaflet phosphorus atoms. The lipid headgroups lie roughly in the region −6Å to 0 (yellow boxes), and the centre of the bilayer is at −20Å (not shown). The location of the molecules in the membrane at the free energy minima are shown in the boxes (red frame for POPC, green frame for POPG; headgroups are shown as red spheres, lipid tails as silver lines, additives as CPK-colored sticks).

**Figure 4.**
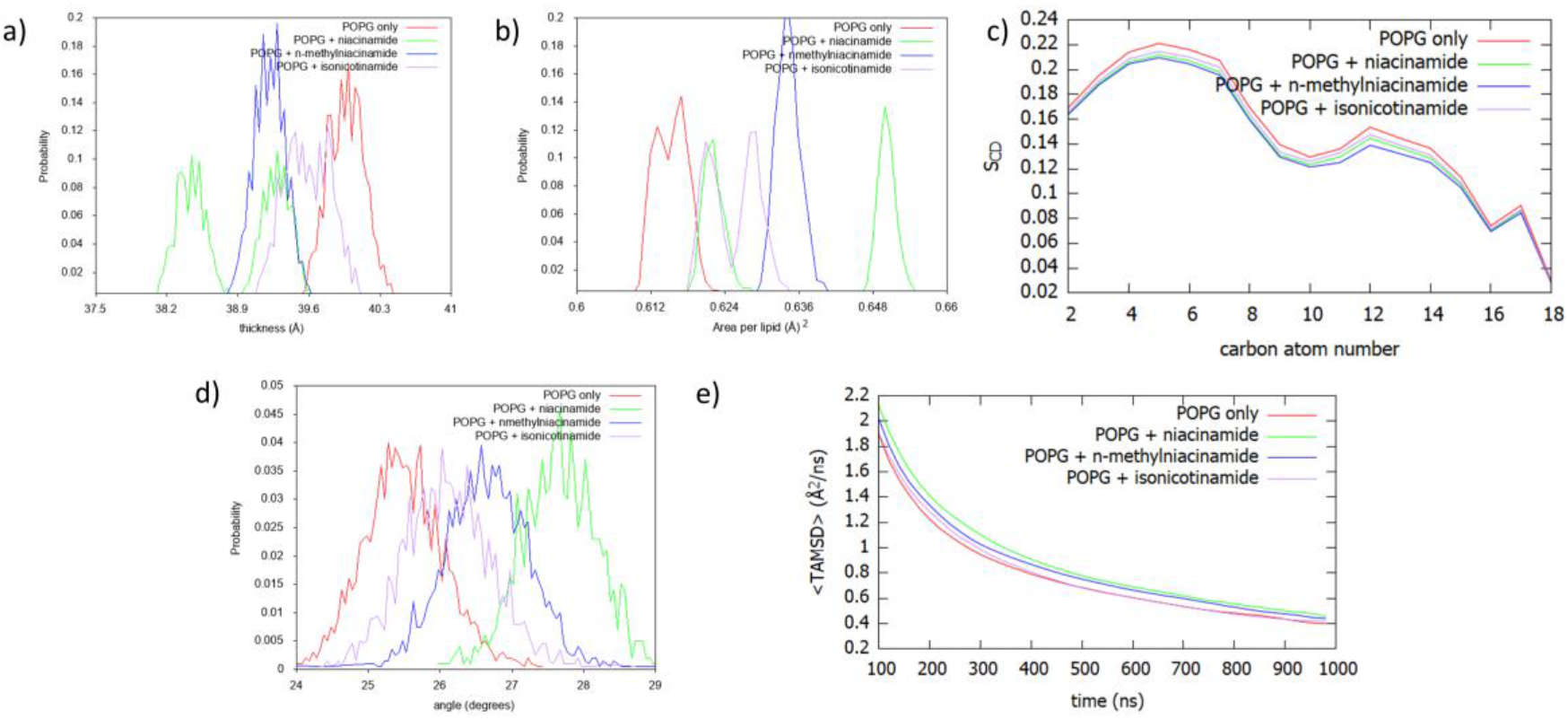
Analysis of unbiased simulations for niacinamide and its derivatives in complex with POPG membranes: a) Distribution of instantaneous membrane thickness values; b) Distribution of area per lipid values; c) S_CD_ order parameter; d) Distribution of lipid tilt angles; e) POPG time-averaged MSD.

**Figure 5.**
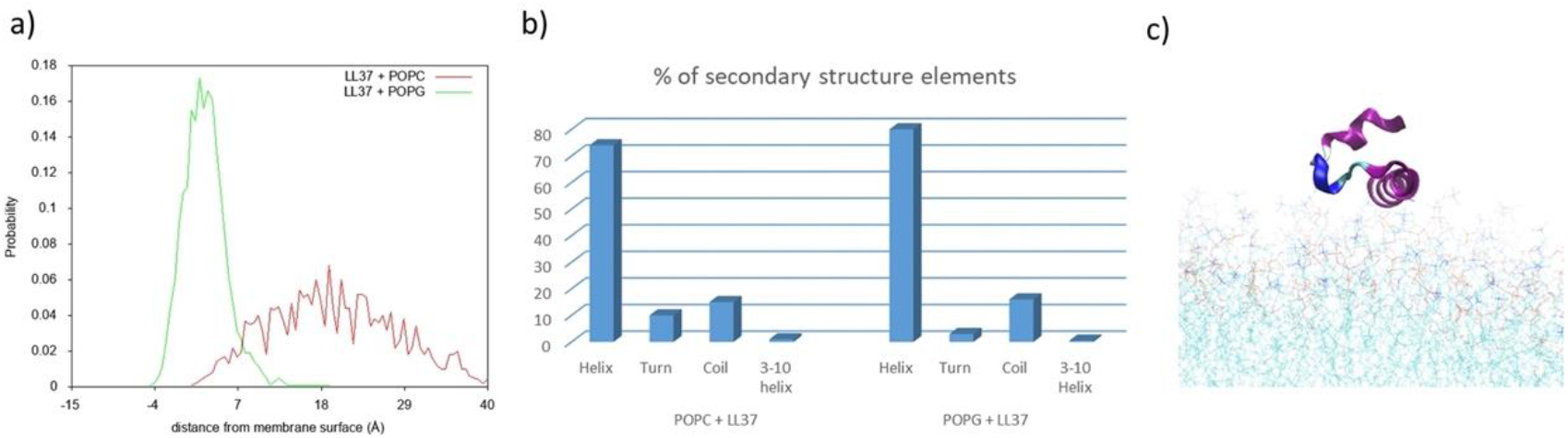
a) Distribution of distances between LL-37 centre of mass and POPG/POPC membrane surfaces. b) Percentage of secondary structure elements in the LL-37 structure in complex with POPG or POPC. c) Snapshot of LL-37 in complex with POPC (purple = alpha helix; blue = coil).

**Figure 6.**
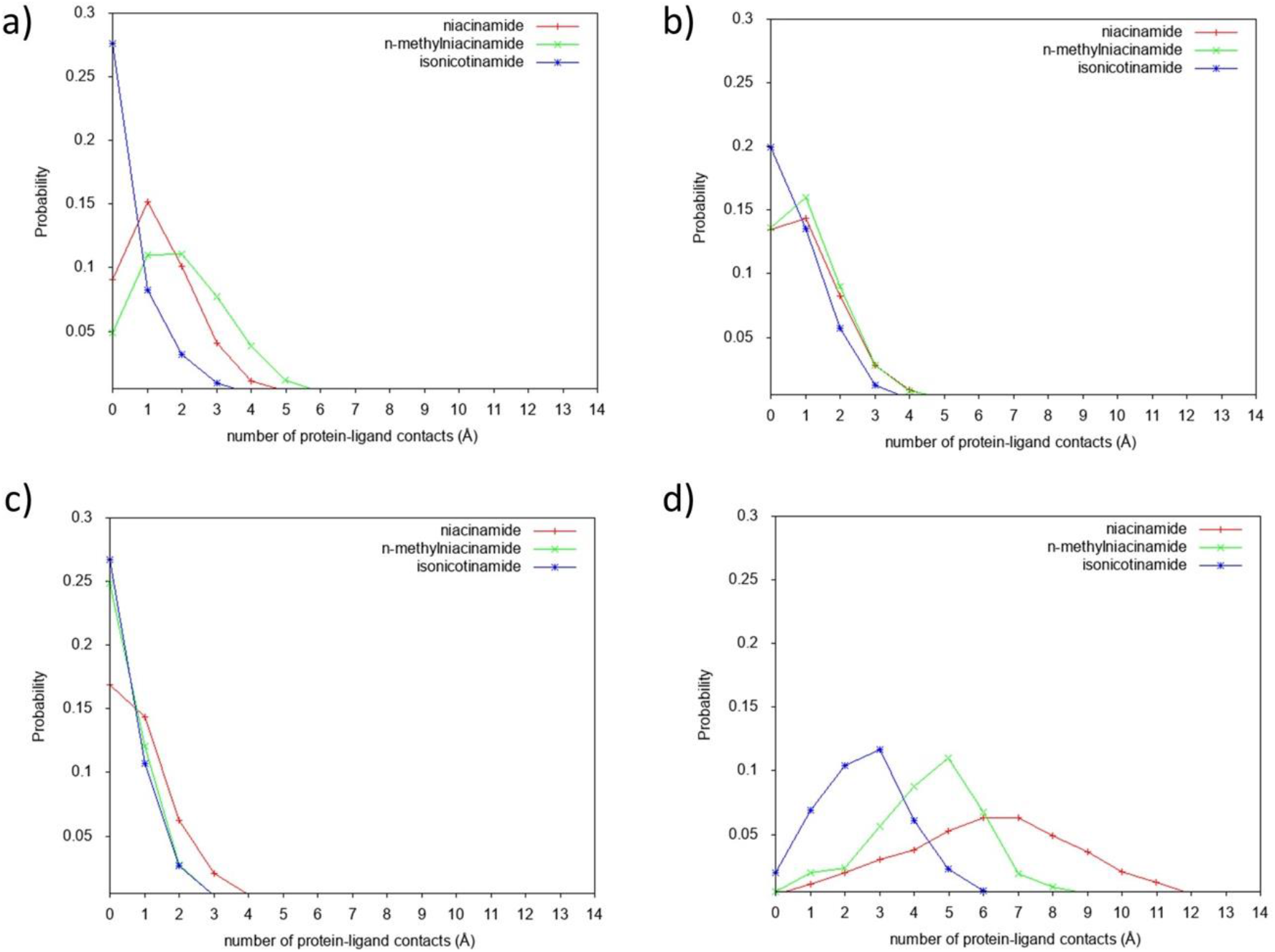
Distribution of contacts for niacinamide and its derivatives with LL-37 in membrane (a) and in three different solvents, water (b), methanol (c) and octanol (d).

**Figure 7.**
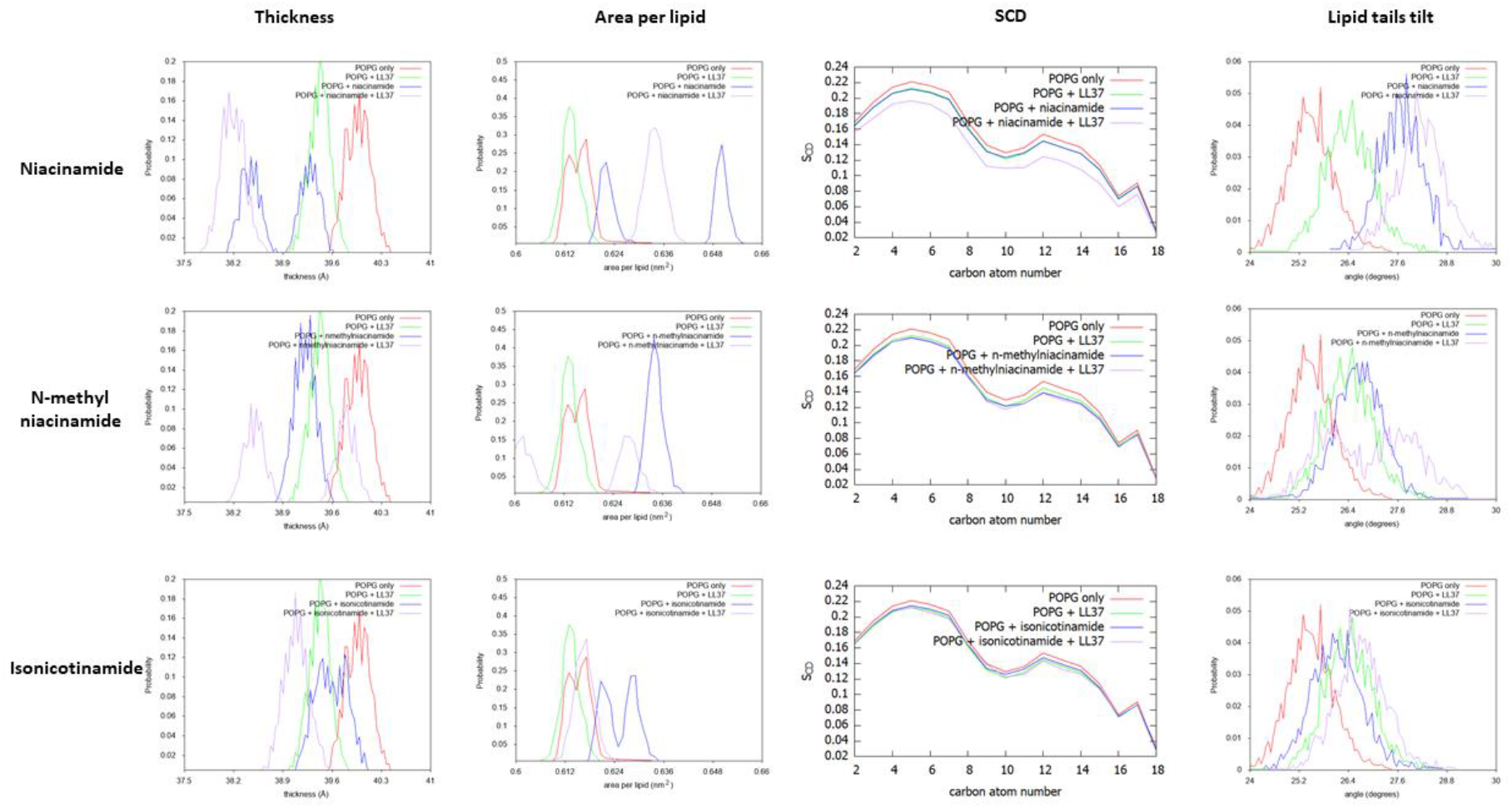
Effect on the membrane structure of the three additives in combination with LL37, as measured by membrane thickness, area per lipid, S_CD order parameter, and lipid tilt angle.

### Microdilution Assay

An overnight Tryptic Soy Agar (TSA) plate culture of *Staphylococcus aureus* ATCC 6538 was scraped and resuspended in 10 mM Sodium phosphate buffer (pH 7.0 ± 0.2) to obtain a bacterial inoculum with cell number of 1 × 10^8^ to 5 × 10^8^ colony forming units per ml (cfu/ml). The inoculum was diluted to 1-5 × 10^6^ cfu/ ml before the assay. The assays were carried out in 96-well TC plates with a final volume of 300 μl. The additives (niacinamide, isonicotinamide and N-methylnicotinamide) and AMP (LL37) and Sodium phosphate buffer (100 mM pH 7.0 ± 0.2, final 10 mM) were added to the wells and the total volume was made up to 165 μl with sterile MilliQ water. One hundred and thirty-five μl of diluted bacterial inoculum was added to the wells, mixed gently, and incubated at 37 ± 0.1°C for 4 h. After the incubation period, aliquots were taken from the reaction mixtures and added to Dey-Engley (D/E) neutralizer broth. The neutralized samples were further diluted, plated onto TSA, and incubated for a minimum 24 h at 37°C. The viable bacteria form colonies on TSA plates after the incubation period that were counted to calculate the recovery [30].

### Molecular dynamics simulations

We ran two independent 1 microsecond-long molecular dynamics (MD) simulations for 22 different systems. Of these, 18 systems consisted of one of the three additives in either a membrane environment or in solution. For the membrane simulations, 20 additive molecules were initially placed on a grid across a 10 × 10 nm patch of POPC or POPG membrane, positioned 5 Å below the membrane surface, corresponding to the average z coordinates of the upper leaflet phosphorus atoms. These lipid-small molecule systems were simulated both on their own, to assess the impact of the additives on the membrane, and in presence of one LL-37 peptide. The peptide was added after 500 ns equilibration and placed in a parallel orientation to the membrane, just above the membrane surface. For the solution simulations, 20 additive molecules were placed in a solvent box of volume ~1.3*10^6^ Å^3^ containing water, methanol or octanol, and with one LL-37 peptide placed in the centre of the box. The remaining 4 simulations were controls consisting of POPC or POPG membranes on their own or containing one LL-37 peptide, but in absence of additives. Table 1 summarises all the simulations performed and lists also the 6 systems used for free energy calculations (see next paragraph).

**Table 1.**
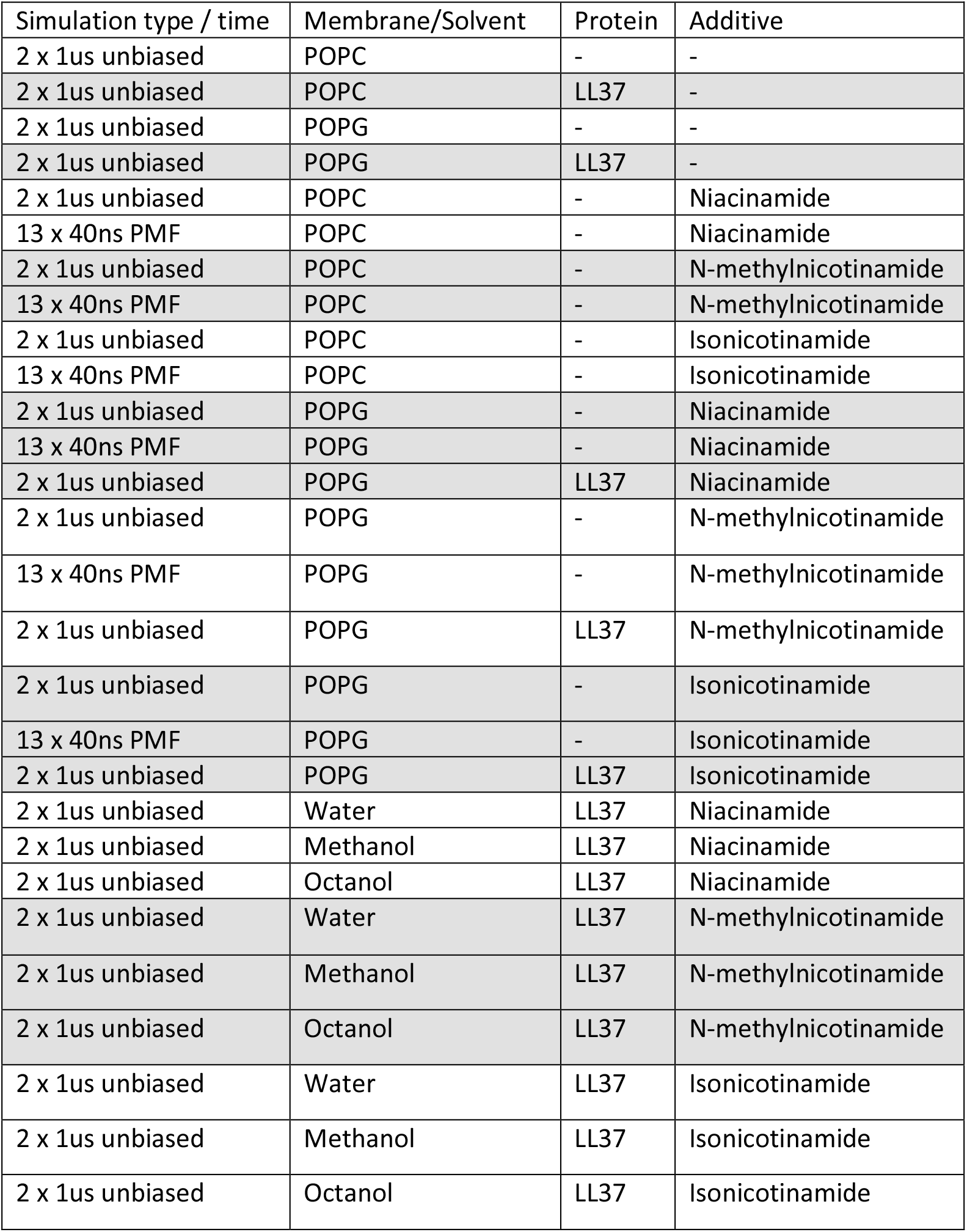
Summary of the systems studied.

| Simulation type / time | Membrane/Solvent | Protein | Additive |
| --- | --- | --- | --- |
| 2 x 1us unbiased | POPC | - | - |
| 2 x 1us unbiased | POPC | LL37 | - |
| 2 x 1us unbiased | POPG | - | - |
| 2 x 1us unbiased | POPG | LL37 | - |
| 2 x 1us unbiased | POPC | - | Niacinamide |
| 13 x 40ns PMF | POPC | - | Niacinamide |
| 2 x 1us unbiased | POPC | - | N-methylnicotinamide |
| 13 x 40ns PMF | POPC | - | N-methylnicotinamide |
| 2 x 1us unbiased | POPC | - | Isonicotinamide |
| 13 x 40ns PMF | POPC | - | Isonicotinamide |
| 2 x 1us unbiased | POPG | - | Niacinamide |
| 13 x 40ns PMF | POPG | - | Niacinamide |
| 2 x 1us unbiased | POPG | LL37 | Niacinamide |
| 2 x 1us unbiased | POPG | - | N-methylnicotinamide |
| 13 x 40ns PMF | POPG | - | N-methylnicotinamide |
| 2 x 1us unbiased | POPG | LL37 | N-methylnicotinamide |
| 2 x 1us unbiased | POPG | - | Isonicotinamide |
| 13 x 40ns PMF | POPG | - | Isonicotinamide |
| 2 x 1us unbiased | POPG | LL37 | Isonicotinamide |
| 2 x 1us unbiased | Water | LL37 | Niacinamide |
| 2 x 1us unbiased | Methanol | LL37 | Niacinamide |
| 2 x 1us unbiased | Octanol | LL37 | Niacinamide |
| 2 x 1us unbiased | Water | LL37 | N-methylnicotinamide |
| 2 x 1us unbiased | Methanol | LL37 | N-methylnicotinamide |
| 2 x 1us unbiased | Octanol | LL37 | N-methylnicotinamide |
| 2 x 1us unbiased | Water | LL37 | Isonicotinamide |
| 2 x 1us unbiased | Methanol | LL37 | Isonicotinamide |
| 2 x 1us unbiased | Octanol | LL37 | Isonicotinamide |

For simulations in solutions, we used the TIP3P model [31] for water and methanol and the CGenFF force field [32] for methanol and octanol. POPC and POPG membranes, composed of 294 and 324 lipids respectively, were described using the CHARMM36 force field for lipids [33]. The structure of LL37 was taken from Protein Data Bank (ID 2K6O) and parameterised with CHARMM27 [34]. Small molecules were generated using the ACEDRG program within the CCP4 suite [35] Fand then parameterised with GAFF force field [36] and AM1-BCC charges [37] through the Antechamber module in AMBER. Membranes were solvated with a 20 Å water layer on each side. Sodium counter ions were used for charge neutralization in water and membrane simulations. Systems in membrane were equilibrated using the following protocol: a) 5000 minimization steps; b) 10 ns with harmonic constraints (1 kcal/mol/A^2^) on protein and lipid heads; c) 10 ns with harmonic constraints (1 kcal/mol/A^2^) on protein only; d) 10 ns without constraints. Systems in solutions were equilibrated by following steps a) and c). Two independent replicas per system were simulated in production runs for 1 microsecond with constant temperature and pressure. All the simulations were carried out with the NAMD 2.9 software [38]

For the simulations in membrane, we analysed the following properties: a) instantaneous membrane thickness measured between the average phosphorus atom z-coordinate for the upper and lower leaflets; b) instantaneous area per lipid; c) deuterium order parameter S_CD_, a parameter typically derived in NMR experiments which reflects the orientational mobility of each C-H bond along the aliphatic lipid tails and thus membrane fluidity; d) the average tilt angle of all lipids with respect to the membrane normal; e) the time average of the mean square displacement (MSD) of lipid molecules, a measure of lipid lateral mobility [25]. All these properties were computed with the MEMBPLUGIN tool [39] For each system, data from the two replica simulations were combined, and the plots show a combined distribution or an appropriate average.

For all simulations including LL37 (in membrane or in solution), we analysed the hydrogen bonds between small molecules and the protein, using as cutoffs 3.5 Å for donor-acceptor distance and 30 degrees for donor-hydrogen-acceptor angle, and their transient unspecific contacts, defined with a maximum distance of 3 Å between any atom of the potentiator and any atom of the protein.

### Free energy estimation

Potential of mean force (PMF) calculations were used to estimate the free energy barrier for each small molecule to penetrate the membrane, by using the adaptive biasing forces (ABF) method as implemented in NAMD [40]. The reaction coordinate was chosen to be the distance between the center of mass (COM) of the potentiator and the surface of the membrane, defined as the instantaneous average of z coordinates of phosphorus atoms of the upper leaflet. The calculations start from the endpoint of the equilibration phase and the reaction coordinate runs from the equilibrated position of the molecules to 10 Å below the membrane surface.

The reaction coordinate was broken down into consecutive windows of size 1 Å, and each one of these was simulated for 40 ns with a force constant of 10 (kcal/mol)/Å^2^ to confine the sampling within the window. The standard error in the PMF was estimated by splitting the calculated data into two block sets of 20 ns per window.

## Data Availability

Simulation data will be made available on the Zenodo platform

## Supporting information

This article contains supporting information.

## Funding and additional information

This work was supported by the STFC Hartree Centre’s Innovation Return on Research program, funded by the Department for Business, Energy and Industrial Strategy.

## Abbreviations and nomenclature

AMP: antimicrobial peptide
PMF: potential of mean force
POPC: 1-palmitoyl-2-oleoyl-sn-glycero-3-phosphocholine
POPG: 1-palmitoyl-2-oleoyl-sn-glycero-3-phospho-(1′-rac-glycerol)
TAMSD: time-averaged mean square displacement
NIA: Niacinamide
MeNAM: N-methylnicotinamide
ISON: Isonicotinamide

